# Comparative analysis of the molecular mechanism of heat shock proteins from pathogenic and non-pathogenic *E.coli*

**DOI:** 10.1101/2020.06.17.157248

**Authors:** Sanchari Bhattacharjee, Mohana Saha, Rakhi Dasgupta, Angshuman Bagchi

## Abstract

Cells can withstand the effects of temperature stress by activating small heat shock proteins IbpA and IbpB. Lon protease employing Ser679 – Lys722 catalytic dyad proteolyze IbpA and IbpB in their free forms, at physiological temperature i.e. without any temperature stress. However, the proteolytic activity of IbpA and IbpB is affected when the catalytic dyad residue of Lon protease is mutated. The mutation S679A in Lon protease brings about some changes so that the proteolytic interactions between the small heat shock proteins with that of the mutant Lon protease are lost which makes a difference in the interaction pattern of mutant Lon protease with their substrates. In the present study, we made an attempt through in-silico approach to figure out the underlying aspects of the interactions between the small heat shock proteins IbpA and IbpB with mutant Lon protease in *Escherichia coli*. We have tried to decipher the molecular details of the mechanism of interaction of proteolytic machinery of small heat shock proteins and mutant Lon protease with S679A mutation at physiological temperature in absence cellular temperature stress. Our study may therefore be helpful to decode the mechanistic details of the correlation with IbpA, IbpB and S679A mutant Lon protease in *E. coli*.

## 1. Introduction

Cells withstand different kinds of stresses while living in a variety of environments. These cellular stresses make the cellular proteins misfolded which affects proper cellular functioning. To maintain proper functioning, the cellular proteins must attain proper folding of misfolded proteins which is a grand challenge of nature. Cell with its own intrinsic anti-stress mechanism counteracts the stressed condition to ensure the proper folding. Cellular stresses are frequent and recurring which makes the cellular proteins to denature, aggregate and damage. The anti-stress response mechanism consists of different types of molecules which elevates at stressed cellular condition such as folding chaperones that ensure the native folding state of the misfolded proteins, Holdase and Disaggregases prevent the misfolding and disaggregation and Proteases eliminate the damaged proteins and thus play a pivotal role in maintaining cellular protein control [1].

Small heat -shock proteins (sHsps) are a type of chaperones with low molecular weight (12-30 kDa) so that are called as molecular chaperones. These proteins protect the stressed cellular protein from further misfolding or aggregation and then assign it to foldase chaperones to regain the native state of the misfolded protein. Two members of small heat shock family, IbpA and IbpB are initially considered as inclusion body protein but later it is observed that they expressed at heat-stressed condition to counter heat-stressed proteins from being misfolded i.e. they play their role as holdase. Both IbpA and IbpB show 48% similarity at amino acid level with a conserved α-crystalin domain flanked N- and C- termini [2].

Lon being the ATP-dependent principal protease in *Escherichia coli* [3] possess a homohexameric structure with a protease and an ATPase domain. Homohexamers of Lon create an internal chamber which triggers degradation damaged proteins [4]. It is reported that at physiological temperature in absence of cellular stress IbpA and IbpB remain in their free form and serves as substrates of Lon protease which depict an interrelation between sHsps and protein degradation machinery [3]. It is observed that, Lon uses a unique Ser^679^ – Lys^722^ catalytic dyad in active site within the protease domain to execute catalysis [5]. This catalytic pocket of Lon protease with the unique Ser-Lys dyad residues plays an important role in substrate binding as well as proteolytic degradation. It is observed and reported that, within the proteolytic domain, Lon protease has the classical catalytic triad residues H665, H667 and D676. But site directed mutagenesis of Lon protease containing substitutions in these classical catalytic triad residues i.e. D676N, H665Y, and H667Y impair the ATPase activity of the enzyme [6]. So the classical catalytic triad residues have no role in proteolysis. Surprisingly, it is also observed that within the proteolytic domain, Lon protease uses a unique conserved Ser679 – Lys722 dyad in active site to promote catalysis [5].This catalytic pocket of the protease domain with Ser-Lys dyad plays a pivotal role in substrate binding. According to an experimental analysis, it is reported that if the catalytic dyad residues Lysine722 and Serine 679 are mutated to Glutamine and Alanine respectively by site directed mutagenesis, then these mutations affect proteolytic activity of Lon protease.

In our previous work, the mode of interactions between IbpA, IbpB and Lon protease is observed at physiological as well as cold and heat-shock temperature and it is observed that Lon interacts more positively with IbpAB at physiological temperature, than those of the stressed temperatures. In the present work, a comparison at physiological temperature has been made through *in-silico* approach to identify the pattern of interactions between IbpA and IbpB with wild type Lon protease as well as mutant Lon protease (S679A) in both pathogenic and non-pathogenic *E.coli* strains. to conclude with a plausible model of molecular mechanism between small heat shock proteins and protein degradation system by Lon protease. So this is the first report to compare and predict the mode of interaction schemes between these proteins in *Escherichia coli* through *in-silico* approach.

## 2. Materials and Methods

### 2.1 Sequence Homology Search and Modeling of the proteins

The amino acid sequences of IbpA, IbpB and Lon protease (UniProt Accession ID: P0C054, P0C058, P0A9M0 and A0A152UUM3 respectively) and their basic information such as sequence length, molecular mass, amino acid modifications were retrieved from UniProt (http://www.uniprot.org/) [7]. In case of UniProt ID P0A9M0, the expression organism of Lon protease is *Escherichia coli* K-12 strain which is a non pathogenic strain whereas in case of UniProt ID A0A152UUM3, the expression organism of Lon protease is *Escherichia coli* O157:H7 strain which is a pathogenic one.

The sequences were used to search for suitable templates in Protein Data Bank (PDB)(www.rcsb.org/) [8] using NCBI PSI-BLAST search tool [9]. For scoring parameters BLOSUM62 matrix is used with the gap cost Existence: 11 Extension: 1 and threshold value is 0.005. In case of both the IbpA and IbpB, the sequence similarity of the obtained PDB templates from BLAST search are less than 30 percent as there was no crystal structure of IbpA and IbpB of *Escherichia coli* in PDB. So, the FASTA sequences of IbpA and IbpB are deposited in RaptorX (raptorx.uchicago.edu/) [10] modelling tools which generated automated models of the proteins. In case of IbpA, RaptorX selected the Heat shock protein 16.0 from *Schizosaccharomyces pombe* (PDB ID 3W1Z) [11] and for IbpB, RaptorX selected the crystal structure of eukaryotic small heat shock protein (PDB ID 1GME) [12] as best template from the database with the highest value of significance. These modelling servers are able to build the entire model of these proteins. The generated models of IbpA and IbpB by RaptorX show an uGDT score of 89 and 88 respectively, a p-value of 1.17e-06 and1.44e-06 respectively with no disordered regions. The modelled protein with high p-value and uGDT score is likely to be a good quality model, so the parameter values of both the modelled proteins indicate that the generated models are with good qualities.

To authenticate the generated model secondary structure prediction is also done using PSIPRED secondary structure prediction server (http://bioinf.cs.ucl.ac.uk/psipred/). For the proteolytic activity, the α-crystallin domain residues are important because Lon protease recognizes this α-crystallin domain and promote proteolysis. It is predicted by PSIPRED Version 3.3 [13] that the α-crystallin domain of IbpA and IbpB spans from amino acid residue 35 to amino acid residue 120 with seven β-sheets and the generated model is also showed the similar secondary structure. So the generated model significantly matches with the predicted structure [Figure 1.A&B].

**Figure 1.**
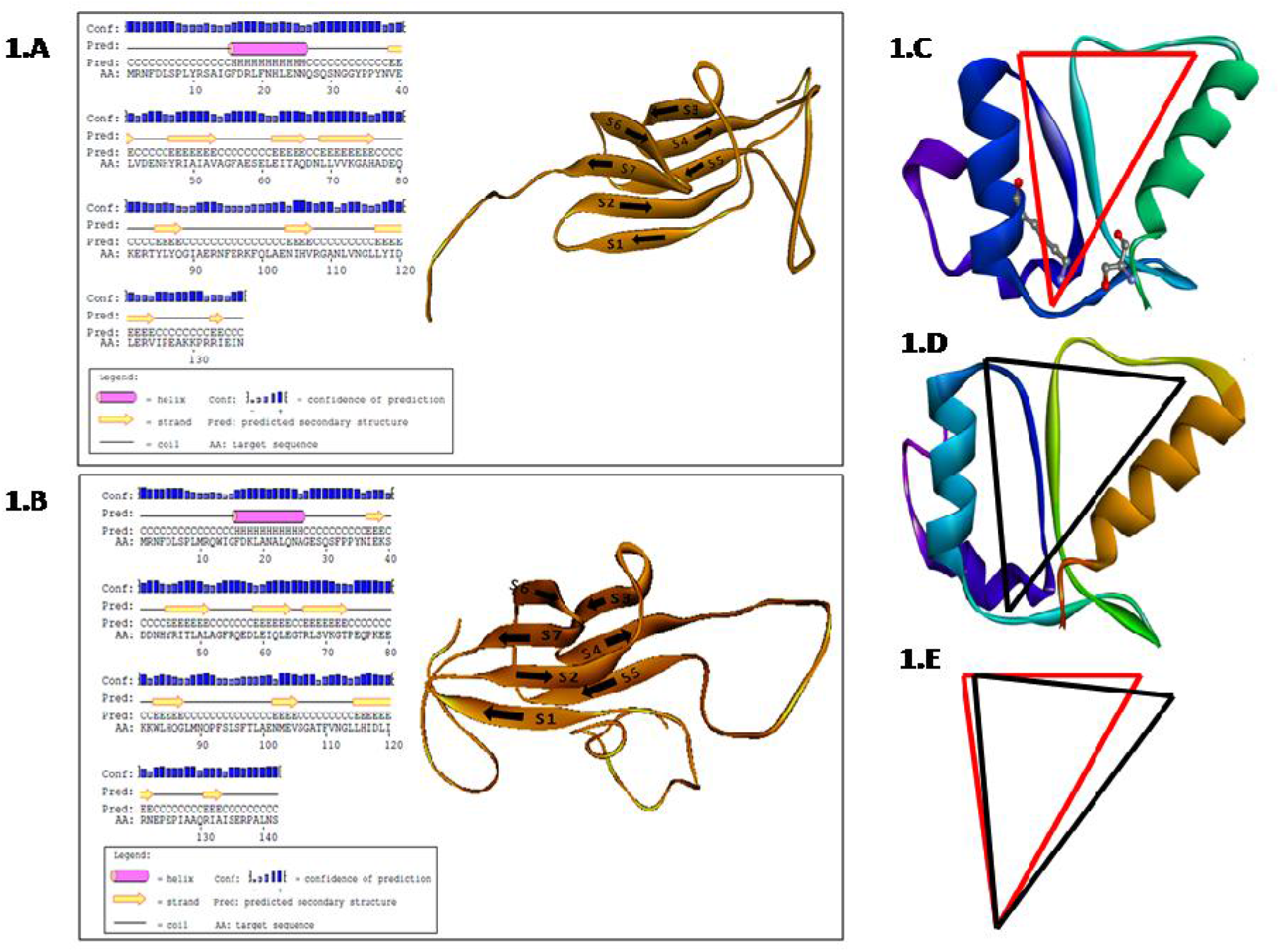
Predicted secondary structure of [A].IbpA and [B].IbpB by PSIPRED version 3.3 (α helix is marked as pink, β strand as yellow, coil as black line; blue bars denote the confidence of prediction). α- crystalline domain (from residue 35 to residue 120) of IbpA and IbpB are composed of seven β-sheets (marked as S1 to S7). [C] Catalytic pocket of wild type Lon protease (marked in red). [D] Catalytic pocket of mutant (S679A) Lon protease (marked in black). [E] Superimposition of catalytic groove of wild type and mutant type Lon protease.

In case of Lon protease, both the FASTA sequences retrieved from non-pathogenic *E.coli* strain (UniProt Accession ID P0A9M0) and pathogenic *E.coli* strain (UniProt Accession ID A0A152UUM3) were used to search for suitable templates in Protein Data Bank (PDB) (www.rcsb.org/) [7] using NCBI PSI-BLAST search tool [8]. For scoring parameters, BLOSUM62 matrix was used with the gap cost: Existence: 11 Extension: 1 with a threshold value of 0.005. In case of both the sequences, the A chain of PDB ID 6U5Z [14] showed the highest query coverage and percentage identity. In case of Uniprot ID P0A9M0 i.e. Lon protease of non pathogenic *E.coli* strain, PDB ID 6U5Z would show 100 % query coverage, 99.87% percentage identity and an E value of 0.0 whereas in case of Uniprot ID A0A152UUM3 i.e. Lon protease of pathogenic *E.coli* strain, PDB ID 6U5Z would show 100 % query coverage, 99.74% percentage identity and an E value of 0.0. Homology modelling of these proteins were performed using SWISS-MODEL (https://swissmodel.expasy.org/) [15] homology modelling server using this PDB ID 6U5Z as a template to generate template based models of the proteins.

The stereo-chemical qualities and the conformations of the resultant structures of IbpA,IbpB and Lon protease were checked with PROCHECK [16] and Verify3D [17] through SAVES server at http://nihserver.mbi.ucla.edu/SAVES/ and Ramachandran plots [18] were drawn. There were no residues present in the disallowed regions of Ramachandran plots. The Verify3D score of IbpA, IbpB and Lon protease would pass the value of 1D-3D profile.

### 2.2 Pairwise Sequence Alignment (PSA) of Lon protease of *E.coli* K-12 strain and Lon protease of *E.coli* O157:H7 strain

FASTA sequences of Lon proteases of *E.coli* K-12 and *E.coli* O157:H7 strains were submitted to Clustal Omega (https://www.ebi.ac.uk/Tools/msa/clustalo/) [19] to perform Pairwise Sequence Alignment (PSA).

### 2.3 Site-directed mutagenesis and secondary structure superimposition of Lon protease

It is reported that if the catalytic dyad residue Serine679 is mutated to Alanine i.e. S679A, this mutation leads to the loss of proteolytic activity [20]. So, site-directed mutagenesis was performed in Discovery Studio version 2.5 to mutate these two Lon proteases at that particular position. The generated mutated structures were energy minimized and tested with PROCHECK and Verify3D. Then these wild type and mutated structures were superimposed using Superpose server (wishart.biology.ualberta.ca/SuperPose/) [21].

### 2.4 Molecular docking with IbpA, IbpB and Lon protease

The models of IbpA and IbpB were docked using Zdock docking server at (http://zdock.umassmed.edu) [22]. The best 10 docked complexes with best docking score were selected and checked with PROCHECK and Verify3D. The docked complex with lowest energy values was selected as final working model.

Next, the above mentioned docking procedures were again performed to dock the IbpAB docked complex with the crystal structure of wild type as well as mutant Lon protease separately in Z-Dock server. Again top 10 docked complexes with best docking scores are selected and in each of these docked complexes are minimized and the energy values are calculated. The docked complexes were tested using PROCHECK and Verify3D. The docked complex with lowest energy scores is selected as final docked complex (IbpAB-Lon protease) [Table 1] [Table 2].

**Table 1:** The best 10 docking poses of IbpA, IbpB and wild type Lon protease showing all energy profiles, PROCHECK and Verify 3D score. The selected docked structure is marked in bold.

| Docked complex | Interaction Energy (IE) (Kcal/Mol) | Van-der-Wal Energy (VDW) (Kcal/Mol) | Electrostatic Energy (EIE) (Kcal/Mol) | Binding Free Energy (BFE) (Kcal/Mol) | RAMPAGE Score (% of disallowed region) | Verify 3D Score |
| --- | --- | --- | --- | --- | --- | --- |
| Complex 1 | -163.36001 | -23.92609 | -139.43392 | -158.1531 | 0% | 89.83% |
| Complex 2 | -157.59543 | -25.32017 | -132.27527 | -152.8694 | 0% | 82.74% |
| Complex 3 | -146.95611 | -20.41863 | -126.53749 | -145.135 | 0% | 94.57% |
| Complex 4 | -136.38073 | -21.26018 | -115.12054 | -144.0759 | 0% | 90.21% |
| Complex 5 | -137.56429 | -25.04836 | -112.51593 | -143.432 | 0% | 90.98% |
| Complex 6 | -131.06914 | -22.30468 | -108.76446 | -144.6112 | 0% | 82.62% |
| Complex 7 | -152.20318 | -23.26713 | -128.93605 | -154.5237 | 0% | 87.92% |
| Complex 8 | -146.59749 | -17.5625 | -129.03499 | -147.2263 | 0% | 88.78% |
| Complex 9 | -161.39836 | -22.13509 | -139.26328 | -153.1479 | 0% | 96.90% |
| **Complex10** | **-172.71306** | **-23.37996** | **-149.3331** | **-164.0855** | **0%** | **98.95%** |

**Table 2:** The best 10 docking poses of IbpA, IbpB and mutant Lon protease showing all energy profiles, PROCHECK and Verify 3D score. The selected docked structure is marked in bold.

| Docked complex | Interaction Energy (IE) (Kcal/Mol) | Van-der-Wal Energy (VDW) (Kcal/Mol) | Electrostatic Energy (EIE) (Kcal/Mol) | Binding Free Energy (BFE) (Kcal/Mol) | RAMPAGE Score (% of disallowed region) | Verify 3D Score |
| --- | --- | --- | --- | --- | --- | --- |
| Complex 1 | -87.07811 | -67.68525 | -19.38986 | -114.0491 | 0% | 94.24% |
| Complex 2 | -141.93129 | -80.60985 | -61.32144 | -152.3293 | 0% | 93.19% |
| Complex 3 | 1316.27711 | 1366.98106 | -50.70395 | 1321.4310 | 0% | 92.67% |
| Complex 4 | -60.10451 | -40.27554 | -19.82897 | -68.5817 | 0% | 93.19% |
| Complex 5 | -114.02465 | -87.57776 | -26.44689 | -126.2604 | 0% | 93.19% |
| Complex 6 | 19794.33187 | 19812.49892 | -18.16706 | 19779.4255 | 0% | 100% |
| Complex 7 | 3926.60519 | 3968.08510 | -41.47991 | 3934.1060 | 0% | 100% |
| Complex 8 | -150.098 | -111.811 | -38.287 | -113.5368 | 0% | 92.67% |
| **Complex 9** | **-190.37358** | **-73.69234** | **-116.68124** | **-215.9242** | **0%** | **94.24%** |
| Complex10 | -97.57960 | 69.80257 | -167.38218 | -85.3113 | 0% | 92.15% |

### 2.5 Molecular Dynamics Simulation of the IbpA-IbpB-Lon protease docked complex

In total, two systems (wild type docked complex and mutant type docked complex) were prepared for simulation. All MD simulations were performed in GROMACS 4.6.5 [23] applying Charmm27 force field [24]. Then, these structures were solvated by explicit solvent SPC/E system [25] water model in cubic boxes with a minimum 10Å distance from the edge of the cube. To obtain a neutral simulation cell the water molecules were replaced by the addition of counter ions (Na^+^ and Cl^−^). Each system was then minimized using steepest descent. The minimized docked complex was again tested with PROCHECK and Verify3D.

A 100 ps NVT equilibration was performed at 310 K with position restraints applied to all of the backbone atoms in order to relieve any bad contacts at the side chain solvent interface. Then position restraints were withdrawn and a 100 ps NPT simulation was conducted. Then 100ns production MD was performed. Snapshots of the trajectory were taken in every 2 ps.

In our previous study, it is reported that, Lon protease interacts with its substrates IbpA and IbpB more efficiently at physiological temperature (310K). In order to study the interaction pattern, both the docked complexes (i.e. wild type Lon-IbpAB and mutant Lon-IbpAB) were performed 100ns of production run in GROMACS version 4.6.5, keeping the pressure constant. The changes in conformation were recorded and plotted against time. Further analyses were done by VMD version 1.9.2 [26] and DS suite. MS Excel was used to generate the graph and for structure visualization, DS Visualizer was used. Relative solvent accessibility (RSA) values are calculated using the XSSP server (http://www.cmbi.ru.nl/xssp/) [27] and superimposition is done using the SuperPose server (wishart.biology.ualberta.ca/SuperPose/)[21]. Finally, these docked complexes were tested using PROCHECK and Verify3D.

### 2.6 System specification

All the work was done using windows operating system and the simulation was performed by Dell Workstation with 2×10 core xenon CPU, 32GB SCC RAM(4X8), 2X2 TB HDD, 1×5GB Nvidia Tesla K20C, 1X1 GB Nvidia Quadro K600.

## 3. Result and Discussions

### 3.1 Sequence similarity and RMSD changes in secondary structure

Pairwise Sequence Alignment reveals that amino acid sequences of Lon protease of *E.coli* non pathogenic K-12 and pathogenic O157:H7 strains are highly conserved [Supplymentary file 1: Pairwise Sequence alignment of Lon protease of *E.coli* pathogenic O157:H7 strain and *E.coli* non pathogenic K-12 strain]. The Percentage Identity Matrix of these two sequences is as follows [Table 3]:-

**Table 3:** Percentage Identity Matrix of Lon protease of *E.coli* pathogenic O157:H7 strain and *E.coli* non pathogenic K-12 strain.

|  |  |  |
| --- | --- | --- |
| tr\|A0A152UUM3\|A0A152UUM3_ECO57 | 100.00 | 99.87 |
| sp\|P0A9M0\|LON_ECOLI | 99.87 | 100.00 |

Pairwise sequence alignment shows that Lon protease is highly conserved whether it is present in pathogenic strain or in non pathogenic strain of *E.coli*. So, the secondary structures of Lon protease of the pathogenic variants and non-pathogenic variants would not have much significant differences as revealed by their backbone RMSD values whereas the Wild type and S679A mutant type Lon protease of the same strain can produce much more significant RMSD values [Table 4]. So, changes in the RMSD values clearly depicts that effect of S679A mutation in the catalytic pocket of Lon protease to have a significant effect on the overall RMSD value and causes more significant RMSD changes in secondary structure irrespective of pathogenicity or non-pathogenicity of the strain.

**Table 4:** RMSD values of superimposed secondary structures of Wild type and S679A Mutant type Lon protease of Non-pathogenic (K-12 strain) and pathogenic (O157:H7 strain) variants of *E.coli*

|  | Wild type Lon<br>(Pathogenic O157:H7 strain) | S679A Mutant type Lon<br>(Pathogenic O157:H7 strain) |
| --- | --- | --- |
| Wild type Lon<br>(Non-pathogenic K-12 strain) | 1.63 Å | 3.21 Å |
| S679A Mutant type Lon<br>(Non-pathogenic K-12 strain) | 3.52 Å | 1.58 Å |

As the effect of S679A point mutation in the catalytic pocket of Lon protease imposes more significant effect in comparison to the pathogenicity or non-pathogenicity of the organism, so the remaining study is focussed on the wild type and S679A mutant type variants of *E.coli* K-12 strain.

### 3.2 Changes in catalytic pocket

In spite of having the catalytic triad residues (His665, His667 and Asp676) Lon protease does not form the catalytic triad rather it employs a unique Ser^679^ - Lys^722^ catalytic dyad for proteolysis. This catalytic dyad residues form a catalytic pocket with a unique structural fold to exert proteolysis. As one of the catalytic dyad residue is mutated (Ser 679Ala), a significant change is shown in the mutated catalytic pocket with respect to the wild type catalytic pocket.

The total SASA (Solvent Accessible Surface Area) value of the catalytic dyad of wild type Lon is 5840.7Å whereas that of the mutant type catalytic dyad is 5912.3 Å. The SASA value is calculated to point out local structural perturbations due to this point mutation for each position of catalytic pocket before and after the mutation.RSA value analysis [Supplymentary file 2: RSA value analysis of catalytic dyad] shows that there is a tendency of transition of core residues to surface residues in the catalytic pocket of mutant type Lon and due to this transition of solvent accessibility, the shape of the catalytic pocket is changed making it wider in the mutant Lon protease.

Superimposition of wild type catalytic pocket and mutant type catalytic pocket generates an overall RMSD change of 1.39Å which considered as a significant change. So, the SASA value, RSA values and RMSD values of these wild type and mutant type catalytic pocket depicts that the mutation in one of the catalytic dyad residue (Ser679Ala) of Lon forms a change in shape in the catalytic pocket of mutant Lon protease which may affect the substrate binding to catalytic pocket as well as the proteolytic activity [Figure 1.C-E].

### 3.3 Changes in binding pose

Binding site analysis showed a change in binding pose of wild type complex and mutant type complex [Figure 2.A & 2.B]. In wild type complex, Lon interacts with IbpB much more than IbpA whereas in case of mutant type complex, Lon interacts with IbpA and IbpB almost equally. Lon actually recognizes these two proteins equally but processes IbpB much faster than IbpA. The tails of IbpA inhibits Lon degradation, so Lon recognizes both these IbpA and IbpB equally but interacts and processes IbpB more as IbpB tails do not inhibit degradation. So, in wild type, Lon interacts with IbpB more and initiates proteolytic degradation. But in case of mutant type, as a result of catalytic dyad residue mutation (S679A), mutant Lon being unable to proteolyse affects the interaction of Lon protease with IbpA and IbpB.

**Figure 2.**
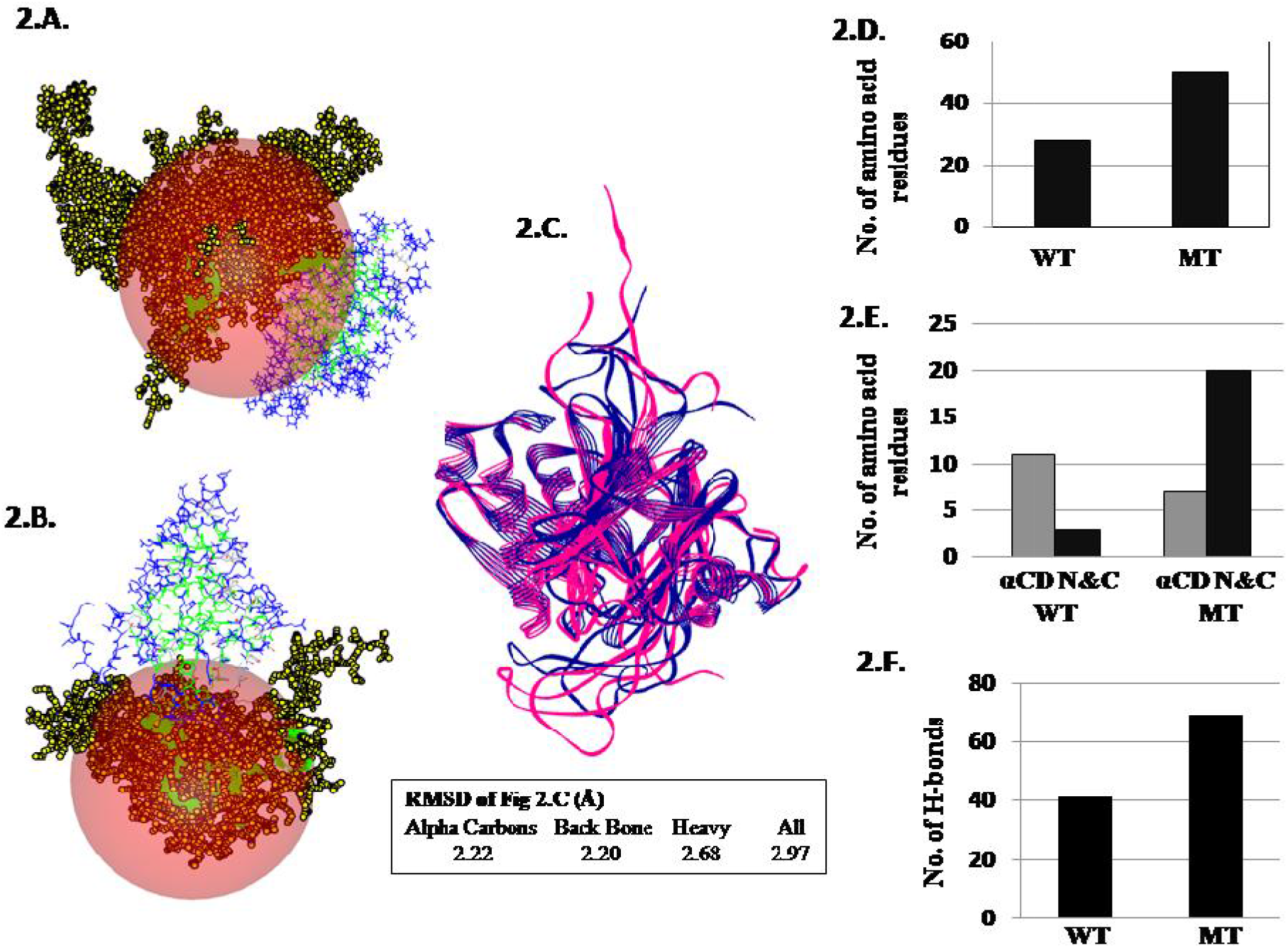
Binding pose of [A] Wild type (WT) and [B] mutant (MT) complexess. Docked IbpA and IbpB are marked in black and Lon protease is in blue and green. Globular pink region indicates the binding surface. [C] Superimposition of WT and MT (IbpAB-Lon). Bar graphs showing [D] No. of amino acid residues involved in binding. [E] Domain specific binding residues of IbpA & IbpB (α-crystalin domain [αCD] and N & C terminal [N&C]). [F] No. of hydrogen bonds involved in WT & MT.

Superimposition of wild type and mutant type docked complex generates an overall RMSD change of 2.97Å which considered as a significant RMSD change. So, these binding pose analysis and RMSD value calculation clearly depicts that the catalytic dyad residue mutation (S679A) of Lon protease affects the involvement of Lon protease with IbpA and IbpB and causes a binding pose alteration in mutant type complex in comparison to wild type complex [Figure 2.C].

### 3.4 Changes in pattern of interaction

#### 3.4.1 Changes in the binding site

Lon protease interacts with IbpA & IbpB in their free form i.e. at physiological temperature whereas the N and C terminal residues of IbpA & IbpB are involved in oligomerization and chaperone activity in temperature stress. Lon recognizes the conserved α-crystalin domain of IbpA & IbpB to promote proteolysis. Binding site analysis revealed that in mutant type, more amino acid residues are involved in binding in corresponding to the wild type. Domain specific binding site analysis showed that in case of wild type complex, the α-crystalin domain residues are involved more in number than N and C terminal residues [Figure 2.D & 2.E] which clearly depicts that the wild type Lon protease initiates proteolysis of IbpA and IbpB at physiological temperature by involving α-crystalin domain residues. But in case of mutant type, it is observed that the N and C terminal residues of IbpA and IbpB are interacting more than α-crystalin domain residues. The mutation in one of the catalytic dyad residue (S679A) hampers the proteolytic activity of Lon protease so that Lon recognize less number of α-crystalin domain residues. In absence of such proteolytic degradation pathway, IbpA and IbpB interact with each other and oligomerize involving N and C terminal residues more in numbers.

#### 3.4.2 Changes in the hydrogen bond interaction

The study reveals that there is a change in the pattern of hydrogen bond interactions. Hydrogen bond analysis shows that more numbers of hydrogen bonds are formed in mutant type in corresponding to the wild type [Figure 2.F]. Wild type Lon protease recognizes the α-crystalin domain of IbpA and IbpB to execute proteolysis. The processing of proteolytic degradation gives an unstable interaction with less numbers of hydrogen bonds. While in case of mutant type, due to the mutation (S679A) in catalytic dyad, the proteolytic activity is hampered partially and this makes mutant Lon unable to recognize the α-crystalin domain residues of IbpA & IbpB properly. Due to this improper recognition, mutant Lon cannot execute the proteolytic degradation properly. Probably, IbpA and IbpB in presence of such partial stress cannot degrade proteolytically but on the other hand interact with each other and form oligomers and makes a stable interaction with more numbers of hydrogen bonds.

### 3.5 Changes in the energy profile

A comparison between the energy profiles of the wild type and mutant type complex was carried out. To analyze this, 148 frames of stable conformations along with the structures are retrieved from molecular dynamics trajectories for each of the docked complexes. Then of these 296 docking poses (148 docked complexes for wild type and mutant type each) the binding free energies (BFE), total interaction energies (TIE), Van-der-Waals energies (VDWE) and electrostatic interaction energies (EIE) are calculated in DS suite. The energy values are tabulated and the unpaired t-test is performed. The unpaired one tailed t-test shows that the alterations of these two data sets are significantly different at P_0.001_ [Figure 3]. It is found that the values of energy profiles are significantly lower in mutant type in comparison to the wild type which states that the mutant type complexes have much more stable interactions throughout the molecular dynamics simulations [Table 5].

**Figure 3.**
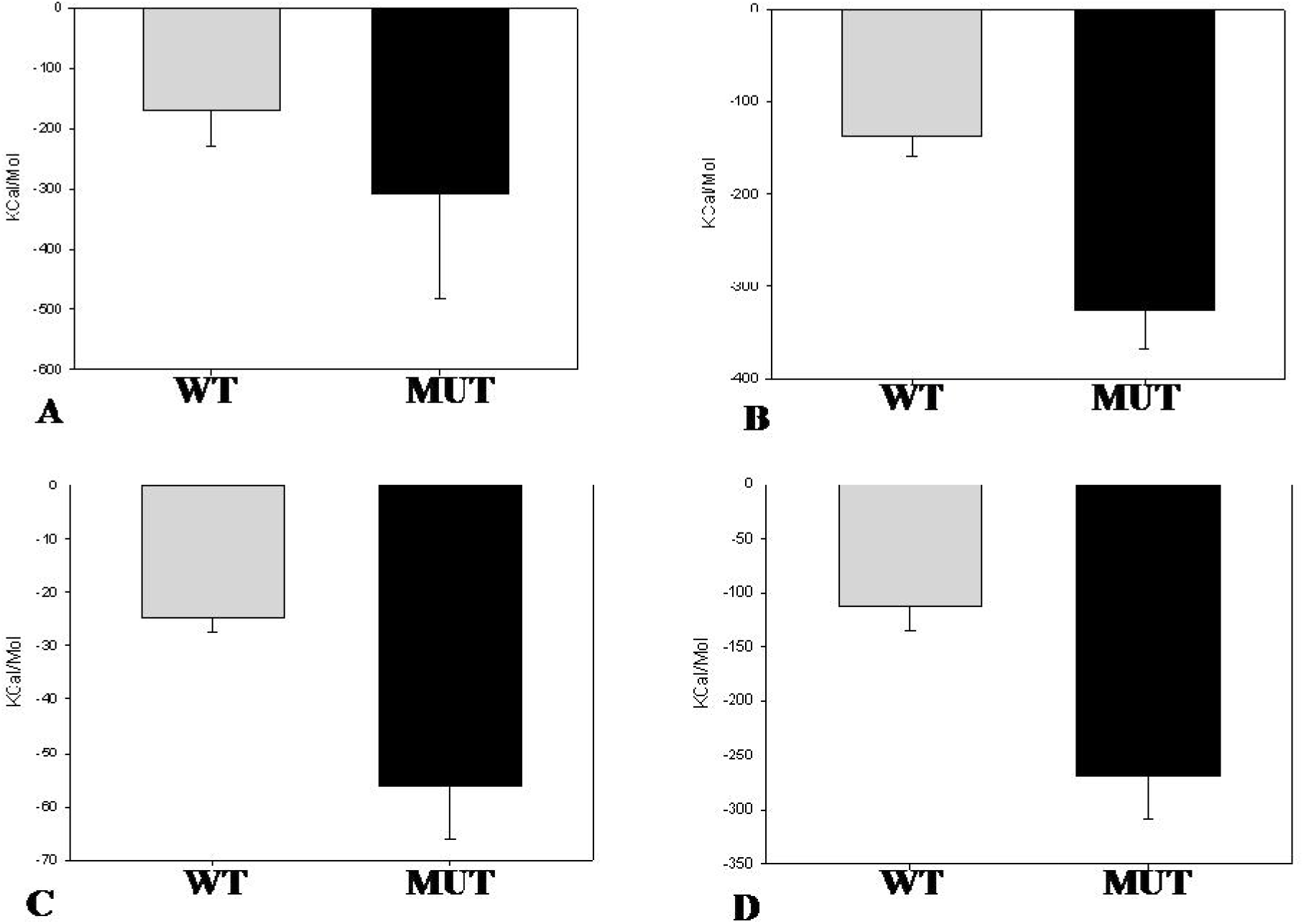
Comparison of interaction energies (Kcal/Mol) (mean±SD) of wild type (WT)(grey bar) and mutated (MUT)(black bar) complexes. **[A]** BFE [**B]** TIE **[C]** VDWE **[D]** EIE. An unpaired one tailed t-test showed all the mutated energy values are significantly less than wild type values (P_0.001_).

**Table 5:**
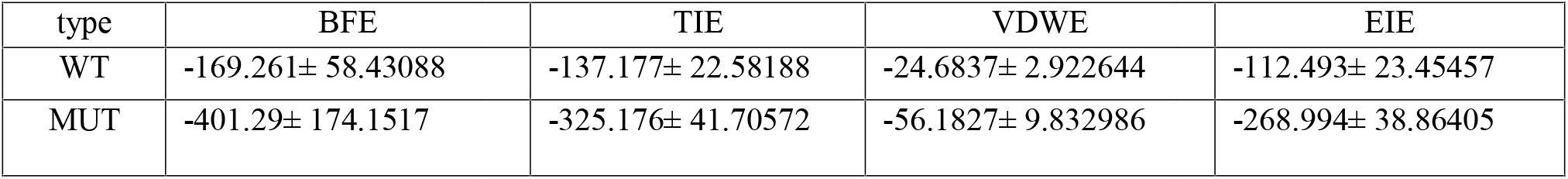
Mean values (Kcal/mol) (mean±SD) at p-value 0.001 and Sample size for all, n = 148 of the energy profiles of wild type and mutant type complexes.

### 3.6 Changes in secondary structure

There also a change is observed in secondary structure of wild type complex and mutant type complex. RMSF values of atoms of wild type and mutant type complexes showed that mean square fluctuation of atoms are much less in mutant type in comparison to the wild type. Less RMS fluctuation points a stable pattern of interaction. In mutant type, due to mutation in one of the catalytic dyad residue (S679A), Lon could not recognize the conserve α-crystalin domain residues of IbpA & IbpB properly and so unable to promote proteolytic degradation properly which in turn gives a stable interaction pattern with less RMS fluctuation.

Secondary structure analysis revealed that the mutant complex confers more structural s bility in corresponding to the wild type complex. Wild type Lon with the help of the catalytic dyad residues exerts proteolytic degradation of IbpA and IbpB which give energetically unfavourable interaction with production of more numbers of coils. On the other hand, the mutant type Lon (S679A) due to it’s mutation in catalytic dyad, cannot degrade IbpA and IbpB proteolytically. In absence of this proteolytic degradation which ought to be exerted by Lon protease, IbpA and IbpB interact and oligomerize with each other forming an energetically favourable and comparatively stable structure with more numbers of α helices, β sheets and turns [Figure 4].

**Figure 4.**
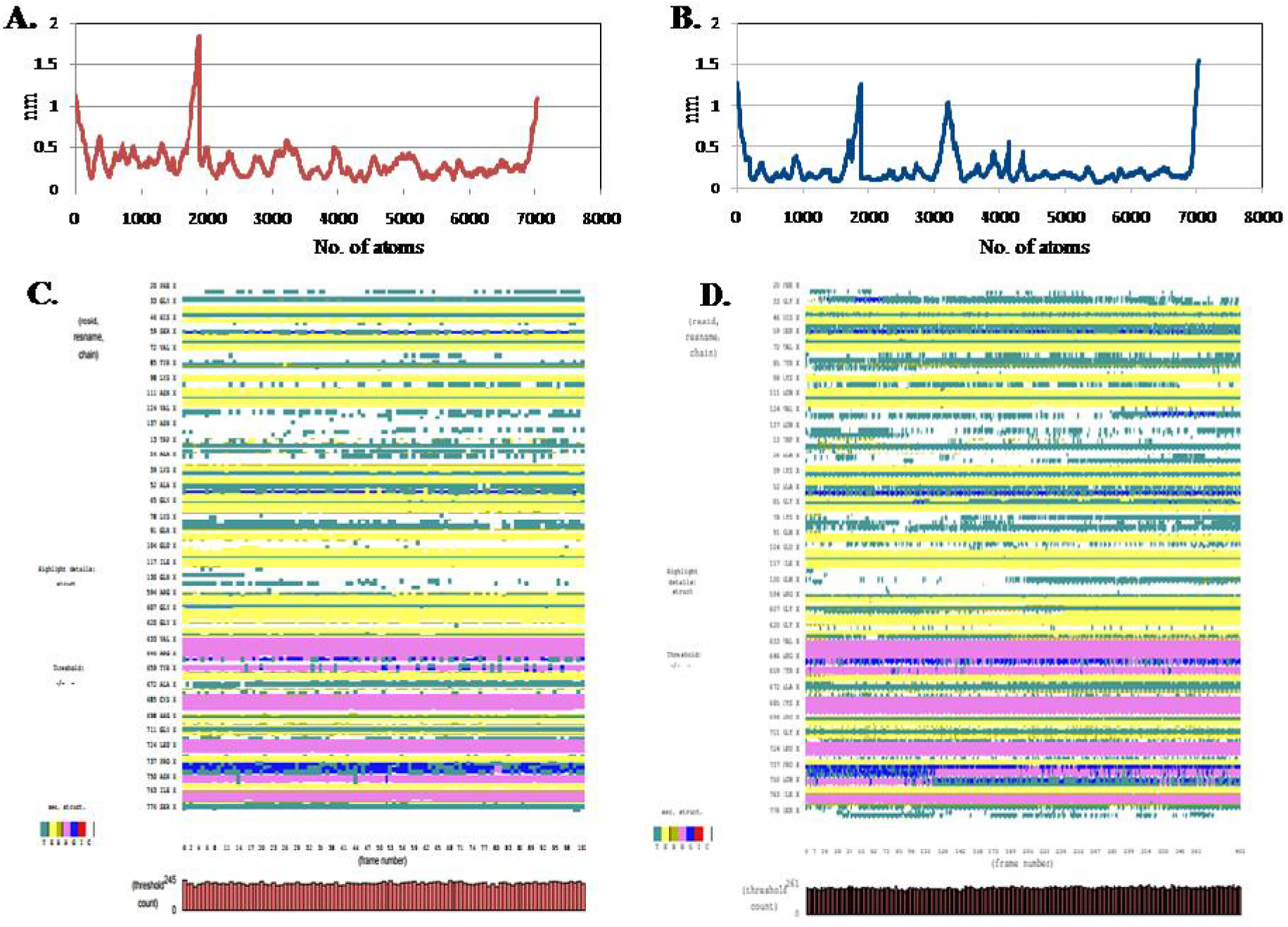
Scatter plot showing atom wise RMSF changes in [A] Wild type complex (red lines) and [B] Mutant type complex (blue lines). Pattern of changes in secondary structure of [C] wild type complex and [D] mutant type complex during simulation where Turn is marked as green, β-sheet as yellow, Alpha helix as pink, 3_10_ helix as blue and Coil as white.

So, the mutant type complex generates more stable secondary structural pattern which clearly depicts the stable interaction pattern of IbpA and IbpB in absence of proteolytic degradation of mutant Lon protease.

So, the mutation in catalytic dyad residue (S679A) in Lon protease makes a difference in the relative solvent accessibility, total solvent accessible surface area that changes the shape of catalytic pocket which in turn affects the substrate binding to Lon protease for initiation of proteolysis. It also causes the alteration of binding pose in comparison to the wild type complex. This mutation also changes the pattern of interaction in mutant type making more stable and favourable interaction throughout the molecular simulation than the wild type which reflects in their hydrogen bond interaction, involvement of binding sites, interaction energy profiles as well as in the values of root mean square fluctuation. Moreover, the secondary structure analysis clearly points that energetically favourable structures are formed more in mutant Lon as there is loss of function due to this mutation. So, S679A mutation in Lon protease generates an overall change in structure and function which makes a change in the interaction pattern of Lon protease with their substrate IbpA and IbpB in *E. coli* even at physiological temperature.

## 4. Conclusion

In this work, we tried to compare the difference in molecular mechanism of pathogenic and non-pathogenic *E.coli* strains. Previously, we made a comparison of the binding interactions in presence of double mutations, viz., S679A and K722Q [28]. However, in this case we did not find much difference between the pathogenic and non-pathogenic *E.coli* strains. Our study is the first of its kind which is based on direct analysis of the heat shock response in both the pathogenic and non-pathogenic *E.coli* strains.

## 5. Acknowledgement

Authors are thankful to the DBT Project No.BT/PR6869/BID/7/417/2012 for financial help and the generous help of Bioinformatics Infrastructural Facility (BIF), University of Kalyani, by providing the necessary machinery requirements and support.

