## Supplementary_Material1 for "Comparative analysis of the molecular mechanism of heat shock proteins from pathogenic and non-pathogenic *E.coli*"

**Supplementary File 1: Pairwise Sequence Alignment of Wild Lon Protease of *E.coli* O157:H7 strain (Pathogenic) and *E.coli* K-12 strain (Non-pathogenic)**

**[The catalytic dyad residue Serine679 and Lysine722 are marked in red colour]**

```
tr|A0A152UUM3|A0A152UUM3_ECO57      MNPERSERIEIPVLPLRDVVVYPHMVIPLFVGREKSIRCLEAAMDHDKKIMLVAQKEAST
sp|P0A9M0|LON_ECOLI                 MNPERSERIEIPVLPLRDVVVYPHMVIPLFVGREKSIRCLEAAMDHDKKIMLVAQKEAST
                                    ************************************************************

tr|A0A152UUM3|A0A152UUM3_ECO57      DEPGVNDLFTVGTVASILQMLKLPDGTVKVLVEGLQRARISALSDNGEHFSAKAEYLESP
sp|P0A9M0|LON_ECOLI                 DEPGVNDLFTVGTVASILQMLKLPDGTVKVLVEGLQRARISALSDNGEHFSAKAEYLESP
                                    ************************************************************

tr|A0A152UUM3|A0A152UUM3_ECO57      TIDEREQEVLVRTAISQFEGYIKLNKKIPPEVLTSLNSIDDPARLADTIAAHMPLKLADK
sp|P0A9M0|LON_ECOLI                 TIDEREQEVLVRTAISQFEGYIKLNKKIPPEVLTSLNSIDDPARLADTIAAHMPLKLADK
                                    ************************************************************

tr|A0A152UUM3|A0A152UUM3_ECO57      QSVLEMSDVNERLEYLMAMMESEIDLLQVEKRIRNRVKKQMEKSQREYYLNEQMKAIQKE
sp|P0A9M0|LON_ECOLI                 QSVLEMSDVNERLEYLMAMMESEIDLLQVEKRIRNRVKKQMEKSQREYYLNEQMKAIQKE
                                    ************************************************************

tr|A0A152UUM3|A0A152UUM3_ECO57      LGEMDDAPDENEALKRKIDAAKMPKEAKEKAEAELQKLKMMSPMSAEATVVRGYIDWMVQ
sp|P0A9M0|LON_ECOLI                 LGEMDDAPDENEALKRKIDAAKMPKEAKEKAEAELQKLKMMSPMSAEATVVRGYIDWMVQ
                                    ************************************************************

tr|A0A152UUM3|A0A152UUM3_ECO57      VPWNARSKVKKDLRQAQEILNTDHYGLERVKDRILEYLAVQSRVNKIKGPILCLVGPPGV
sp|P0A9M0|LON_ECOLI                 VPWNARSKVKKDLRQAQEILDTDHYGLERVKDRILEYLAVQSRVNKIKGPILCLVGPPGV
                                    ********************:***************************************

tr|A0A152UUM3|A0A152UUM3_ECO57      GKTSLGQSIAKATGRKYVRMALGGVRDEAEIRGHRRTYIGSMPGKLIQKMAKVGVKNPLF
sp|P0A9M0|LON_ECOLI                 GKTSLGQSIAKATGRKYVRMALGGVRDEAEIRGHRRTYIGSMPGKLIQKMAKVGVKNPLF
                                    ************************************************************

tr|A0A152UUM3|A0A152UUM3_ECO57      LLDEIDKMSSDMRGDPASALLEVLDPEQNVAFSDHYLEVDYDLSDVMFVATSNSMNIPAP
sp|P0A9M0|LON_ECOLI                 LLDEIDKMSSDMRGDPASALLEVLDPEQNVAFSDHYLEVDYDLSDVMFVATSNSMNIPAP
                                    ************************************************************

tr|A0A152UUM3|A0A152UUM3_ECO57      LLDRMEVIRLSGYTEDEKLNIAKRHLLPKQIERNALKKGELTVDDSAIIGIIRYYTREAG
sp|P0A9M0|LON_ECOLI                 LLDRMEVIRLSGYTEDEKLNIAKRHLLPKQIERNALKKGELTVDDSAIIGIIRYYTREAG
                                    ************************************************************

tr|A0A152UUM3|A0A152UUM3_ECO57      VRGLEREISKLCRKAVKQLLLDKSLKHIEINGDNLHDYLGVQRFDYGRADNENRVGQVTG
sp|P0A9M0|LON_ECOLI                 VRGLEREISKLCRKAVKQLLLDKSLKHIEINGDNLHDYLGVQRFDYGRADNENRVGQVTG
                                    ************************************************************

tr|A0A152UUM3|A0A152UUM3_ECO57      LAWTEVGGDLLTIETACVPGKGKLTYTGSLGEVMQESIQAALTVVRARAEKLGINPDFYE
sp|P0A9M0|LON_ECOLI                 LAWTEVGGDLLTIETACVPGKGKLTYTGSLGEVMQESIQAALTVVRARAEKLGINPDFYE
                                    ************************************************************

tr|A0A152UUM3|A0A152UUM3_ECO57      KRDIHVHVPEGATPKDGPSAGIAMCTALVSCLTGNPVRADVAMTGEITLRGQVLPIGGLK
sp|P0A9M0|LON_ECOLI                 KRDIHVHVPEGATPKDGPSAGIAMCTALVSCLTGNPVRADVAMTGEITLRGQVLPIGGLK
                                    ************************************************************

tr|A0A152UUM3|A0A152UUM3_ECO57      EKLLAAHRGGIKTVLIPFENKRDLEEIPDNVIADLDIHPVKRIEEVLTLALQNEPSGMQV
sp|P0A9M0|LON_ECOLI                 EKLLAAHRGGIKTVLIPFENKRDLEEIPDNVIADLDIHPVKRIEEVLTLALQNEPSGMQV
                                    ************************************************************

tr|A0A152UUM3|A0A152UUM3_ECO57      VTAK	784
sp|P0A9M0|LON_ECOLI                 VTAK	784
                                    ****
```
